# AutoGenome: An AutoML Tool for Genomic Research

**DOI:** 10.1101/842526

**Authors:** Denghui Liu, Chi Xu, Wenjun He, Zhimeng Xu, Wenqi Fu, Lei Zhang, Jie Yang, Guangdun Peng, Dali Han, Xiaolong Bai, Nan Qiao

**Affiliations:** Huawei Technologies Co., Ltd; CAS Key Laboratory of Regenerative Biology and Guangdong Provincial Key Laboratory of Stem Cell and Regenerative Medicine, Guangzhou Institutes of Biomedicine and Health, Chinese Academy of Sciences, Guangzhou, China; Guangzhou Regenerative Medicine and Health Guangdong Laboratory, Guangzhou, China; Key Laboratory of Genomic and Precision Medicine, Beijing Institute of Genomics, Chinese Academy of Sciences, Beijing, China

**Keywords:** AutoGenome, AutoML, Residual fully-connected neural network, Genomic, Deep learning

## Abstract

Deep learning have made great successes in traditional fields like computer vision (CV), natural language processing (NLP) and speech processing. Those achievements greatly inspire researchers in genomic study and make deep learning in genomics a very hot topic. Convolutional neural network (CNN) and recurrent neural network (RNN) are frequently used for genomic sequence prediction problems; multiple layer perception (MLP) and auto-encoders (AE) are frequently used for genomic profiling data like RNA expression data and gene mutation data. Here, we introduce a new neural network architecture, named residual fully-connected neural network (RFCN) and demonstrate its advantage for modeling genomic profiling data. We further incorporate AutoML algorithms and implement AutoGenome, an end-to-end automated genomic deep learning framework. By utilizing the proposed RFCN architectures, automatic hyper-parameter search and neural architecture search algorithms, AutoGenome can train high-performance deep learning models for various kinds of genomic profiling data automatically. To make researchers better understand the trained models, AutoGenome can assess the feature importance and export the most important features for supervised learning tasks, and the representative latent vectors for unsupervised learning tasks. We envision AutoGenome to become a popular tool in genomic studies.

---

In the last decades, the emergence of high throughput sequencing technology revolutionized the biomedical research and generated tons of omics data. In genomics area, microarray^1^ and next generation DNA-Seq^2^ are widely used to identify genome-wide copy number variations, singlenucleotide polymorphism (SNP) and DNA mutations; in epigenomics area, MeDip-Seq^3^, BS-Seq^4^ are used to profile DNA methylations; ChIP-Seq is used to identify chromatin associate proteins^5^; in transcriptomics area, microarray^1^ and RNA-Seq^6^ are used to quantify whole RNA expressions profile; in proteomics area; LC-MS^7^ and ICAT^8^ are used to analyze protein complex and quantify proteins; in metabolomics area, MNR^9^ and mass spectrometry ^10^ are used to profile metabolic markers. Omics data provide comprehensive information at different molecular system levels, and have been widely used in biomedical researches^11,12^, and at the same time numerous bioinformatics tools been developed for analyzing omics data.

Deep learning have been proved to be very effective in areas like computer vision^13,14^, natural language processing^15^ and speech processing^16–18^. By leveraging properly designed, deep, stacked neural network architecture, low/middle/high level features could be extracted automatically and combined together to predict the learning target in an end-to-end fashion. Inspired from the achievements of deep learning in the traditional areas, researchers are now actively designing neural network architectures for genomic study, which makes deep learning in genomics study a very hot topic (Fig 1a).

**Figure 1.**
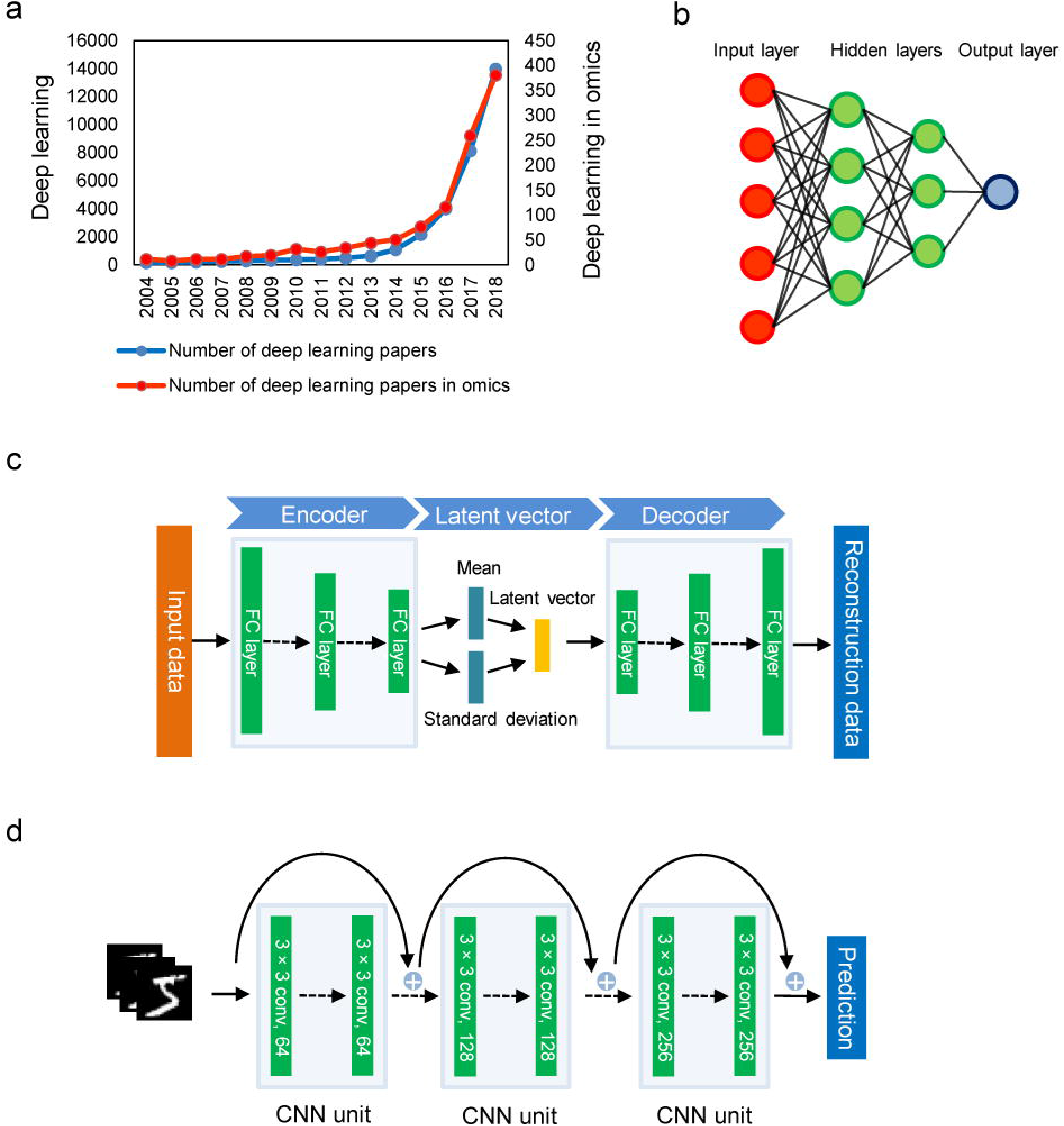
Introduction to deep learning neural network. a). Publication trends of deep learning papers vs. deep learning papers in genomics. Data is from https://apps.webofknowledge.com. b). Multiple layer perception (MLP) architecture. Information from previous layer is used as input for the next layer. c). Variational autoencoder (VAE) architecture. Encoder compresses raw data into latent vectors by decreasing number of neurons in each layer. Decoder reconstructs the raw data from latent vectors. d). Residual convolutional network architecture. Residual network involves skip-connections in the neural networks. The basic unit in the network are convolutional layers.

Convolutional neural network^14^ (CNN) and recurrent neural network^16^ (RNN) are widely used neural network architectures. CNN could effectively extract multiple-dimension spatial features from 1-D to N-D data^14,19,20^, and RNN could capture long short-term features^21^ from time series data. By treating the DNA/RNA/protein sequence as image data or sequence data, CNN and RNN could then be used to model the genomic sequence problems like variant discovery^22^, DNA motif finding^23^, sgRNA on-target site prediction^24,25^, protein-protein-interaction prediction^26,27^, and drug design^28–30^.

However, for non-sequence data like genomic profiling data, where RNA expression value, gene mutation status, or gene copy number variations from whole genome are profiled, CNN or RNN will be invalid because there are no spatial or temporal relationships in the data. For genomic profiling data, the underlying mechanism is the gene regulatory pathway/network: several genes interact with (activate or inhibit) each other and compose a structured hieratical network to regulate the biological functions^31^, which is the key to model genomic profiling data using deep learning.

A lot of studies using multiple layer perception (MLP), autoencoders (AE) or variational autoencoders (VAE) for supervised or unsupervised tasks of genomic profiling data (Fig. 1b-c), and the researchers observed improvements compared with traditional machine learning algorithms^32–34^. Evidences from deep learning studies in computer vision reveal that network depth is crucial important^14,35,36^. However both MLP and VAE face the problem of vanishing gradient^37,38^, which means they cannot train very deep neural network. Most of the published paper in genomics deep learning have less than four hidden layers (Fig. S4a).

Hyper parameter selection and network architecture selection usually take the researchers a big paragraph to discuss, and they are even harder for deep learning beginners. The development of automated machine learning (AutoML) aims to remove the gaps and make AI more democratize to everyone. Automatic model selection, feature selection, hyper-parameter search and neural architecture search are commonly used in AutoML^39–43^. In genomic area, however, most studies are still limited to handcrafted DNN structures and parameter space.

To address those challenges and facilitate genomic studies with deep learning, we build AutoGenome, an end-to-end AutoML framework for genomic study. In AutoGenome, we propose residual fully-connected neural network (RFCN) and its variants, and validate their performance outperforming MLP and VAE. We further adopt hyper-parameter search and efficient network architecture search^41^ (ENAS) algorithm to AutoGenome, to enable it automatically searching for novel neural network architectures. We show that AutoGenome can be easily used to train customized deep learning model, evaluate the model performances, and interpret the results for genomic profiling data.

## Residual fully-connected neural network outperform MLP in genomic profiling data modeling

Looking back to the success of deep learning in ImageNet challenge^44^, we could find that increasing network depths plays an important role^14,35,36^. However, simply increased the network depth would cause the problem of gradient vanishing^37,38^. Residual network was then proposed to resolve this problem. Residual network prevent vanishing gradient by adding a short cut at each layer which makes the gradient backpropagation smoothly (Fig. 1d). ResNets^45^, HighwayNets^46^ and DenseNets^47^ are major variants of residual network. Residual network could also strengthen the feature propagation and encourage feature reuse.

To make the residual network also works for genomic profiling data, we propose residual fully-connected neural network (RFCN). Different from previous residual networks^45,47^, which utilize convolution layer as basic unit, residual fully-connected neural network employs fully-connected layer as the basic unit, each layer might connect to the next layer directly (Fig. 2a left, Path 1), or through a skip connection (Fig. 2a left, Path 2), or through a branch (Fig. 2a left, Path 3). The basic fully-connected (FC) unit is shown in Fig. 2a right. The FC unit comprises of a sequential of fully-connected layer, batch normalization layer and activation function layer.

**Figure 2.**
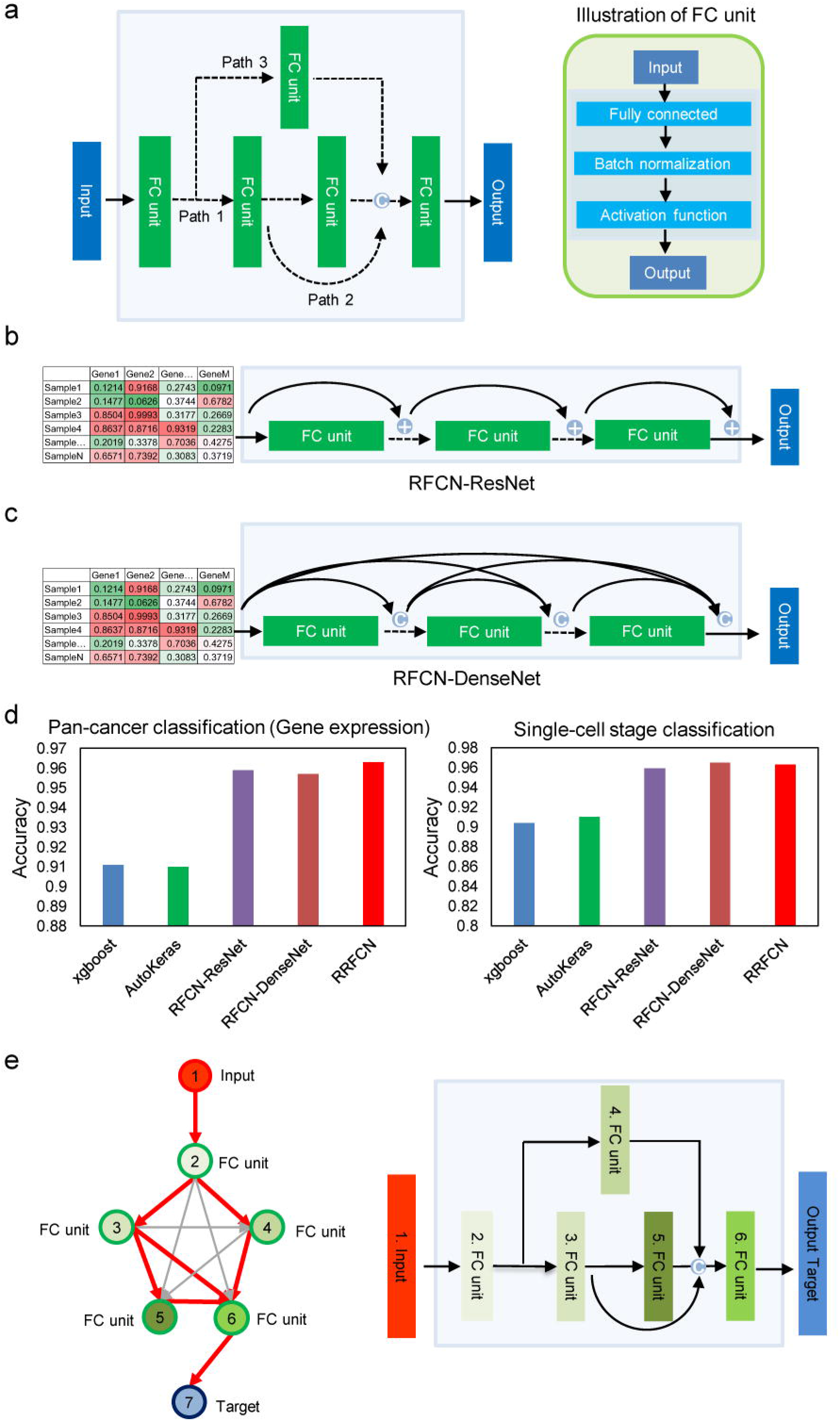
Residual fully-connected neural network (RFCN). a). We introduce fully-connected (FC) units into residual network and name it as RFCN. Each FC unit contains sequential layers of FC, batch normalization and activation function. We construct three kinds of variants for RFCN - b) RFCN-ResNet, c) RFCN-DenseNet and e) randomly-wired residual fully-connected neural network (RRFCN). b) In RFCN-ResNet, for each FC unit, it takes the summation of the output and input of the former unit as its input. “+” indicates the summation operation. c) In RFCN - DenseNet, for each FC unit, it takes the concatenation of all the outputs from previous units as its input. “C” indicate the concatenation operation. d). Performance of the three variants of RFCN compared with traditional methods (XGBoost and AutoKeras) for pan-cancer type classification and single-cell stage classification based on gene expression profiles. The y-axis indicates top-1 classification accuracy. e). In RRFCN, ENAS searches for an optimal subgraph from a large directed acyclic graph (DAG) as the final best model. Arrows indicate all the potential connections among nodes, while red ones indicate the searched optimal subgraph connections (left). A searched optimal subgraph corresponds to a neural network (right).

By replacing the CNN unit with FC unit in the standard form of ResNet/DenseNet, we propose the RFCN variants: RFCN-ResNet and RFCN-DenseNet (Fig. 2b and 2c). In RFCN-ResNet, for each FC unit, it takes the summation of the output and input of the previous unit as its input (Fig. 2b). In RFCN-DenseNet, for each FC unit, it takes the concatenation of all the outputs from previous units as its input (Fig. 2c). To make researchers experience RFCN-ResNet and RFCN-DenseNet easily, we implemented the algorithms in AutoGenome, which could be used to search for the best RFCN-ResNet/RFCN-DenseNet structures and hyper parameters automatically (see Methods).

To assess the performance of RFCN-ResNet/RFCN-DenseNet, we design experiments on two independent datasets and compare the performances with XGBoost^48^ (A popular machine learning algorithm) and AutoKeras^39^ (A popular AutoML tool). In the first experiment, we use gene expression profiles from 8,127 samples, corresponding to 24 tumor types from TCGA^49^ dataset to perform a pan-cancer classification task; somatic mutation profiles from 6,186 TCGA samples (24 tumor types) are also used to perform the pan-cancer classification task independently. In the second experiment, we use the gene expression profile from 10,000 mouse single cells, covering 10 different embryonic developmental stages (downloaded from a public dataset^50^) to perform a classification task. After preprocessing the datasets (Details in Methods), we train different models with XGBoost, AutoKeras, RFCN-ResNet and RFCN-DenseNet separately and evaluate their performances with an independent test dataset. The results shows that for pan-cancer classification task using gene expression profiles, the RFCN-ResNet and RFCN-DenseNet achieve accuracy of 95.9% and 95.7%, and outperform those obtained by using XGBoost and AutoKeras by 4.8% and 4.9%, respectively (Fig. 2c); with only genomic somatic mutation profiles, the RFCN-ResNet and RFCN-DenseNet achieve accuracy of 64.1% and 60.1%, and outperform XGBoost and AutoKeras by 5.7% and 19.1% (Fig. S1a). In the mouse single cell classification experiment, RFCN-ResNet and RFCN-DenseNet achieve accuracy of 95.9% and 96.5%, and outperform those obtained by using XGBoost and AutoKeras, by 6.1% and 5.5%, respectively (Fig. 2d). These results show us that the proposed RFCN-ResNet and RFCN-DenseNet have significantly better performances than the MLP based architectures (from AutoKeras) in genomic profiling data modeling.

## Randomly-wired residual fully-connected neural network is a promising architecture in genomic research

According to the definition of ResNet and DenseNet, the skip-connection structures in RFCN-ResNet and RFCN-DenseNet are fixed. The neural network architectures are still in a hand-crafted manner, which might not be the best choice for various kinds of scientific problems. We therefore propose another RFCN variant named randomly-wired residual fully-connected neural network (RRFCN), which adopt neural architecture search (NAS) to generate and search for the best residual fully-connected neural architecture for any given genomic profiling problems. With RRFCN, researchers could search for brand new RFCN architectures for their scientific problems.

There are a lot NAS optimization methods^40–43^, and we implement the efficient neural architecture search^41^ (ENAS) algorithm into AutoGenome to search for RRFCN. ENAS could greatly accelerate the training efficiency by forcing all the child models to share weights^41^. The RRFCN search space are represented as a single directed acyclic graph (DAG, Fig. 2e left).Therefore, a RRFCN architecture can be realized by taking a subgraph from the DAG (the subgraph is represented as the red path in Fig. 2e left, and the corresponding architecture is shown in Fig. 2e right). The building block for RRFCN is the FC unit illustrated in Fig 2a right, and the search space are show in Methods.

We also assesse the performance of RRFCN on the previous two datasets. For the pan cancer study, the RRFCN achieve the best accuracy compared with Xgboost, AutoKeras, RFCN-ResNet, RFCN-DenseNet for both gene expression and gene mutation data (the accuracy is 96.3% for TCGA gene expression profiles and 68.1% for TCGA somatic mutation profiles, Fig. 2d left, Fig. S1a). The best RRFCN architectures from both dataset have six hidden layers and four skip connections, but the four skip connections connect differently (Fig. 3a, Fig. S1b); both RRFCN architectures are brand new neural network architecture in genomic research. For the mouse single cell experiment, the RRFCN achieve an accuracy of 96.3%, ranked the second highest, slightly lower than RFCN-ResNet (Fig 2d right), the best RRFCN architecture has seven hidden layers and seven skip connections (Fig. S2b), which is also a novel neural network architecture in genomic research.

**Figure 3.**
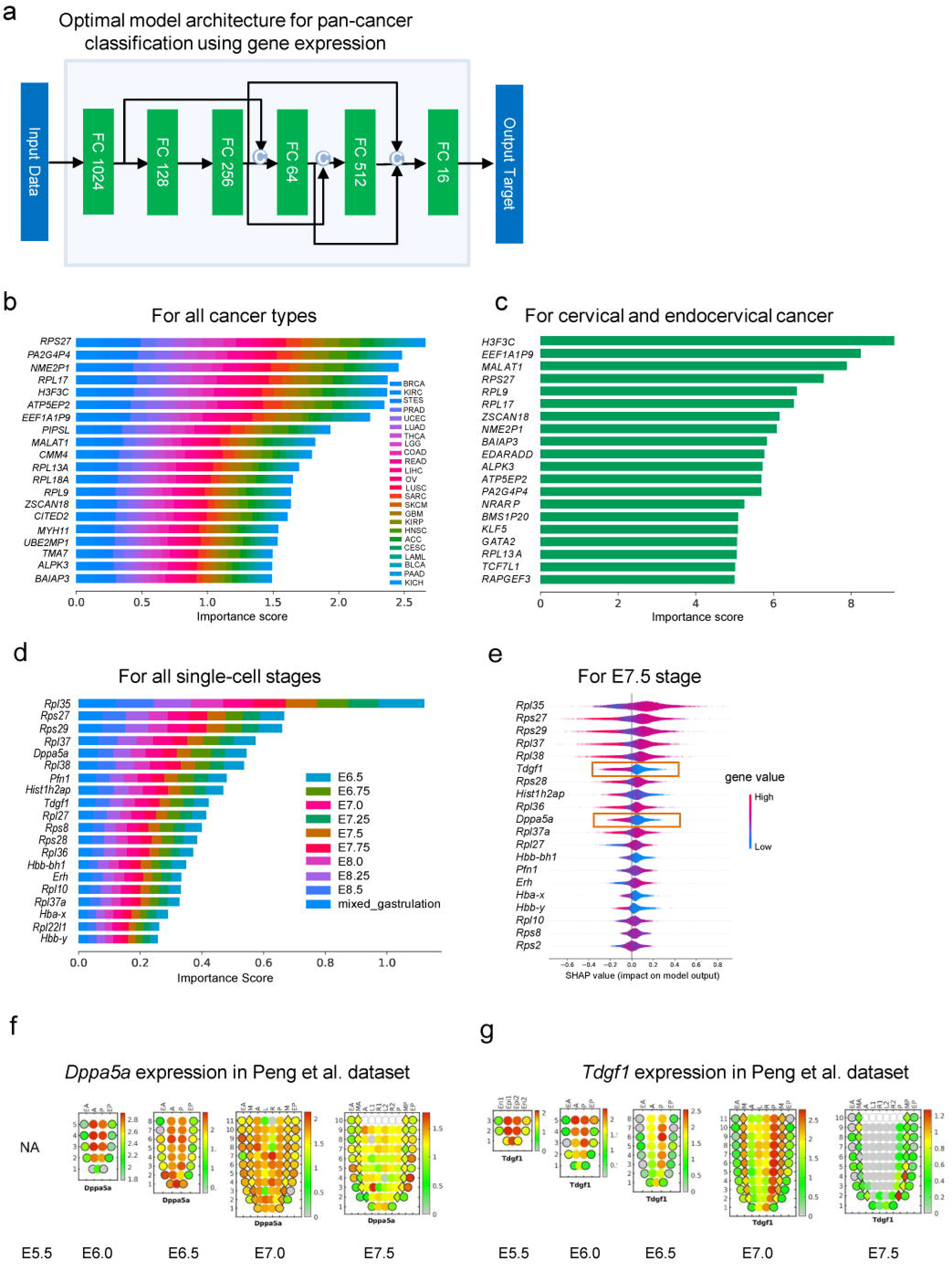
Feature importance analysis for explaining predictions of RFCN. a). The optimal model architecture for pan-cancer classification using gene expression profiles. Top-20 genes ranked by feature importance scores from gene expression based cancer type classification for b) all cancer types and c) a specified-cancer, cervical and endocervical cancer. Top-20 genes ranked by feature importance scores from gene expression based single-cell classification for d) all single-cell stages and e) E7.5 stage. e). For E7.5 stage, high- and low-gene expression are indicated in purple and blue. Red rectangles indicate marker genes for early embryonic development. f). *Dppa5a* and g). *Tdgf1* gene expression from E5.5 to E7.5 stages in Peng et al. dataset.

## Explain predictions for RFCN models

A lot researchers believe deep learning models are black boxes^51–53^, which are difficult to explain. To facilitate the researchers to investigate into the deep learning models, we bring a popular method called SHapley Additive exPlanations^54^ (SHAP) into AutoGenome. Given a deep learning model, SHAP will calculate the marginal contribution for each feature to the overall predictions, which is referred as SHAP value^54^. AutoGenome could visualize the feature importance of each gene to the predicted classes (Fig. 3b, d), or the SHAP values distribution of each gene to the predicted classes (Fig. 3c, e; Fig. S3). Those visualizations provide meaningful insights toward the deep learning models.

For the pan-cancer classification task with gene expression profiles, the top important genes for each cancer type are visualized by AutoGenome (Fig. 3b, c; Fig. S3a, b; Supp. Table 1), and a lot of literature support are found for the genes in the top list. For example, the top 1 rank gene across all 24 cancer types is *RPS27* (Fig. 3b), which has been observed highly expressed in various human cancers^55,56^; the pseudogenes *PA2G4P4* and *H3F3C* are reported to be functional in many cancers^57,58^ (Fig. 3b); in cervical and endocervical cancers (CESC), a long non-coding RNA gene,*MALAT1,* ranks top 3 among all 8449 genes (Fig. 3c), and its high expression predicts a poor prognosis of cervical cancer^59^; in lung adenocarcinoma (LUAD), *ST3GAL5* ranks top 9 (Fig. S3a), and this gene participates in modulation of cell proliferation and maintenance of fibroblast morphology, and is known to be associated to in situ pulmonary adenocarcinoma^60^; in lung squamous cell carcinoma (LUSC), *BMS1P20* ranks top 8 (Fig. S3b), it is known to associate with lung cancer^61^. The full list of top important genes for each cancer type are shown in Supp. Table 1, those results offer potential biomarker candidates for cancers.

We also analyze the top important genes for the somatic-mutation-profile-based pan-cancer classification. *TP53* and *PICK3CA* rank top 1 contribution for the prediction of ovarian serous cystadenocarcinoma (OV) and breast invasive carcinoma (BRCA) respectively, showing positive effect to the model output with positive SHAP values for their mutated status in most samples (Fig. S3c-d). It is consistent with *TP53* and *PICK3CA* being most frequently mutated in OV and BRCA among the 24 cancer types, with frequency of 87.34% and 32.41% respectively (276 OV patients with *TP53* mutations vs. 316 total OV patients; 318 BRCA patients with *PIK3CA* mutation vs. 981 total BRCA patients). Both *TP53* and *PICK3CA* mutations are reported to be potential diagnostic and prognostic biomarkers for ovarian cancer^62^ and breast cancer^63^ respectively.

For the single-cell embryonic developmental stage classification task, the top important genes for each developmental stage are also visualized by AutoGenome (Fig. 3d-e; Supp. Table 2). A lot ribosomal genes are observed in the top list, which is consistent with previous finding that ribosome genes play an important role in embryonic development and stem cell differentiation^64,65^. From the gene SHAP value distributions per developmental stage, researchers could better understand how the genes contribute to the development stages, for example, the top 1 rank gene, *Rpl35* (Fig. 3d-e), has been reported to be important during early embryogenesis^66,67^, and the *Rpl35* SHAP distribution in developmental stage E7.5 (Fig. 3e) shows clearly that high *Rpl35* expression value will predict E7.5 and low *Rpl35* expression value will not predict E7.5; *Dppa5a, Tdgf1* are also on the top list and have been reported to regulate early embryonic development and pluripotency^68–71^. By looking into their SHAP distribution in developmental stage E7.5 (Fig. 3e), it’s also clear that their low expression will predict E7.5 and their high expression will not predict E7.5. The observations are also consistent with another independent dataset, where *Dppa5a* and *Tdgf1* have relatively high expression values in E6.0, E6.5 and E7.0, but drop significantly in E7.5 (Fig. 3f, g).

## Residual fully-connected VAE outperform traditional VAE in omics data

RFCN-ResNet, RFCN-DenseNet and RRFCN implemented in AutoGenome could be used for supervised learning tasks like regression or classifications, we also extend RFCN to unsupervised learning by proposing a new neural network architecture named residual fully-connected variational auto-encoder (Res-VAE, Fig. 4a) by adding skip connections to the traditional variational auto-encoder^72^ (VAE) architecture. In traditional VAE architecture (Fig. 1c), the encoder compress input into latent vectors by decreasing the number of neurons in each subsequent layer, the decoder reconstruct the input data from the latent vectors. Res-VAE add skip-connections to both the encoder and the decoder, to strengthen the feature propagation and encourage feature reuse for each module. By minimize the sum of reconstruction loss and Kullback-Leibler divergence (KLD) loss, Res-VAE learns the representative features from the data, the representative features are stored in the latent vectors and could be used for further analysis^73–75^.

**Figure 4.**
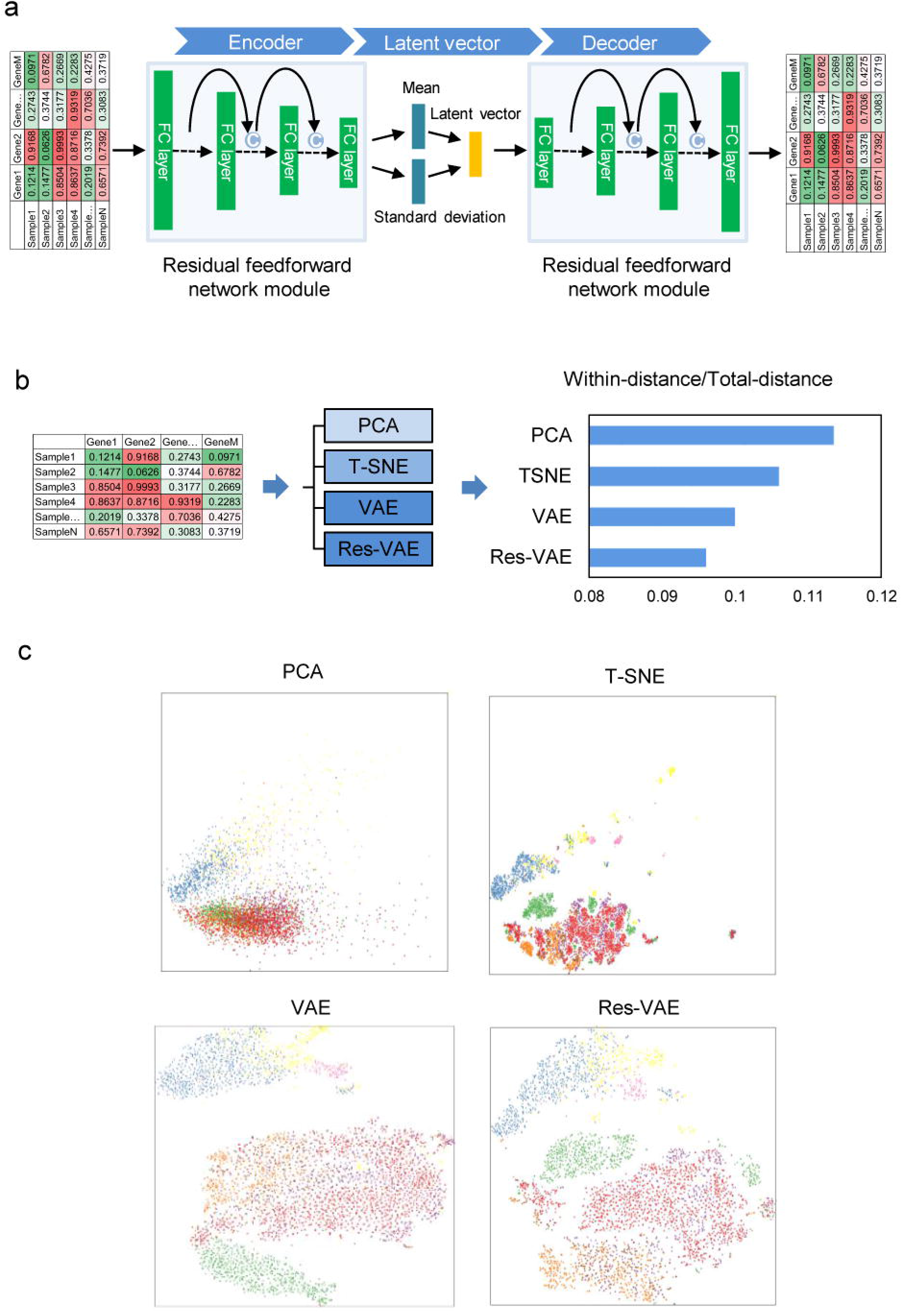
Residually fully-connected VAE. a). Res-VAE compresses the raw data into latent vectors in the Encoder process, and reconstructs the raw data from latent vectors in the Decoder process. Skip-connections are involved both in the Encoder and Decoder parts. “C” indicate the integration methods of information from different layers is concatenation. b). Within-distance/Total-distance between single-cell categories using their gene expression profiles for comparison of Res-VAE and other three unsupervised machine learning methods. c). 2D visualization for single cell categories after processing by Res-VAE and other three methods.

To assess the performance of Res-VAE, we design an experiment using a new single-cell RNA-seq dataset provided by 10X platform^76^ and compare the result of Res-VAE with PCA, T-SNE and VAE (Fig. 4b). In general, the unsupervised learning of PCA, T-SNE, VAE and Res-VAE are trying to mapping the high dimensional gene expression data (16653 genes in total, see Methods) in to a low dimensional space. For fare comparison, the first two dimensions from PCA and T-SNE results are visualized in Fig. 4c. For VAE and Res-VAE, the number of latent variables are set to 128, which means in the encoder module, the expression value of 16653 genes are compressed into 128 variables, T-SNE are then used to map the 128 latent variables into 2 dimensional space, also shown in Fig. 4c. The color in Fig. 4 indicate the cell types, and it’s quite straight forward that Res-VAE results have more clear boundaries between different cell types and tight clusters for each cell type. Davies-Bouldin score^77^ is usually used to measure the clustering goodness, and smaller Davies-Bouldin score indicate better clustering. The results showed Res-VAE have the smallest Davies-Bouldin score, indicating all the cell type clusters are very compact (Fig. 4b).

## AutoGenome - An AutoML tool for Genomic Research

Researchers usually encounter a lot challenges in applying deep learning in their research, for example “ which deep learning framework shall I use”, “ how to prepare the data”, “ how to choose a good model architecture” and “how to set the hyper parameters”. AutoML aim to solve these problems by combining many advanced technologies like automated data clean^78^, automated feature engineering^79^, hyper parameter optimization^80^ and neural architecture search^40–43^. Some AutoML tools have been developed like AutoKeras, Auto-sklearn, H2O AutoML, but currently they are only able to search MLP, CNN or RNN based architectures.

Here we developed a new AutoML tool called AutoGenome for genomic research, to enable researchers to perform end-to-end learning with the best cutting edge neural network architectures easily. When a gene matrix data from genomic profiling is provided, for supervised learning tasks, AutoGenome could automatically search for the best MLP architectures, the proposed RFCN-ResNet architectures, RFCN-DenseNet architectures and RRFCN architectures; after AutoGenome find the best model, the evaluation confusion matrix is also provided; at last, AutoGenome could calculate and visualize the feature importance score and SHAP value distributions, which could be used for the researchers to further investigate and explain the models(Fig. 5a). For unsupervised learning tasks, AutoGenome could automatically search for the best Res-VAE architectures, after AutoGenome find the best model, the latent variable matrix and reconstruction matrix are also provided for further analysis (Fig. 5a).

**Figure 5.**
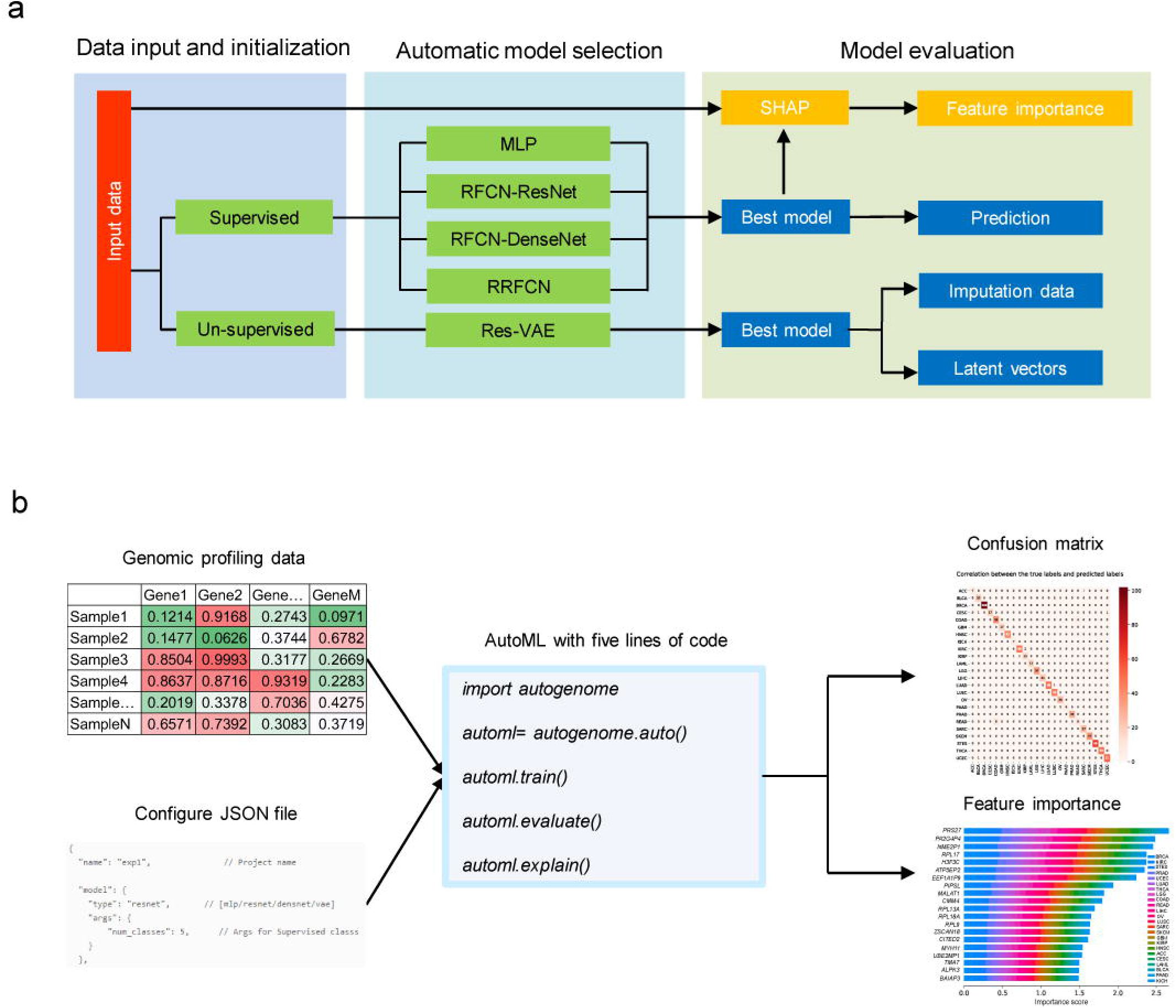
Overview of AutoGenome. a). The process of automatically building a model by AutoGenome includes three steps: data input and initialization (left), automatic model selection (middle) and model evaluation (right). There are two modes of tasks – supervised and unsupervised tasks, and five model structures – MLP, RFCN-ResNet, RFCN-DenseNet, RRFCN and Res-VAE for selection. B) Input data and codes for accomplishing AutoGenome. AutoGenome takes a genomic profiling data matrix and configuration JSON file selecting modes and model structures as input, and outputs model accuracy, confusion matrix and feature importance scores for explaining predictions. All the processes above takes only five lines of code for AutoGenome.

With AutoGenome, researchers only need to execute five lines of code to perform the whole analysis (Fig. 5b). Along with the great convenience, AutoGenome also provide the expert mode for sophisticated deep learning researchers, all the parameters in AutoGenome could be manually changed by modify configure file, you can even change the hyper parameter search space to your favorite sets, the configure file is well organized in JSON format (Fig. 5b).

## Discussion

Genomic profiling data grows rapidly, and they are very important in biomedical researches. Similar to the recent achievements in computer vision, natural language processing and speech processing, novel neural network architectures are needed to overcome the challenges in this area. Compared with the widely used MLP architecture, the advantages of the RFCN architecture are obvious: a. RFCN could be used to train deeper neural network; b. RFCN could strengthen the feature propagation and encourage feature reuse; c. RRFCN could be used to generate novel RFCN architectures. With AutoGenome, it would be easy for researchers to practice RFCN in their research.

After reviewed several publications in genomic research, we found that most of the published MLP based neural network have less than 4 layers (Fig. S4a). We also summarized the depth (number of hidden layers) of the best model for RFCN architectures in experiments 1 and 2 (Supp. Table 3), the depth of RFCN-ResNet and RRFCN are around 6 to 7, the depth of RFCN-DenseNet are around 11, it’s obvious that RFCN architectures tend to train deeper neural networks. We also investigate the correlations between the depth of RRFCN and accuracy (Fig. S4b), and found that the best accuracy is from the model with the depth around 6 to 7, further increasing the depth wouldn’t improve the performance any more. This might be relevant to the complexity of the problems as well as the underlying gene regulatory pathway/network.

In our experiments, RFCN architectures are proved to have better performances than MLP architectures, but for the three RFCN variants (RFCN-ResNet, RFCN-DenseNet and RRFCN), the performances are similar (Fig. 2d). Compared with RRFCN, RFCN-ResNet and RFCN-DenseNet are easier to train and the architectures are easier to understand (Fig. 2a, b, c, e).

AutoGenome is open source and free available for everyone. Notebook examples are also provided to help researchers experience AutoGenome smoothly.

## Methods

### AutoGenome AutoML tool

AutoGenome is built on Tensorflow^81^ and implemented as a Python package. The parameters required for training are specified in the JSON configuration file. It can be saved, revised and loaded to temporary workspace. The JSON configuration file comprises of the following modules. 1) Data loader module. User can specify the path to the training, evaluation and test data set, or set the number of threads, capacity when loading data. 2) Model trainer module. In this module, Users can specify the general training parameters, like batch size, number of GPUs utilization, learning rate range, and so on. 3) Loss function module. User can specify the loss calculation method. Now we support “cross entropy” for classification task, “mean-square error” for regression task, “area under curve” for imbalanced binary classification task. 4) Search mode module. There are five mode that can be selected, MLP, FC-ResNet, FC-DenseNet, ENAS for supervised learning and Res-VAE for unsupervised learning. Each mode have different search space and we will discuss in the following context 5) Best model save path and reload module. User can specify the path to save the best model for sharing or future reloading.

### Hyper-parameter search

Hyper-parameter search method refers to previous described approach^82^. The general hyper-parameters in search space are learning rate, total batch size, momentum, weight decay, number of layers in neural networks and number of neurons in each layers. The specific search space for each mode is follows:

### MLP Search Space

Search space are as followings.1) The number of neuros in each layer, default value is [8, 16, 32, 64, 128, 256, 512, 1024]. 3) The drop-out ratio of the first layer compared with the input layer, selected from [0.6, 0.8, 1.0]

### RFCN-ResNet Search Space

Search space are as followings. 1) The number of blocks for ResNet, default value is [1, 2, 3, 5, 6]. 2) The number of neuros in each layer, default value is [8, 16, 32, 64, 128, 256, 512]. 3) The drop-out ratio of the first layer compared with the input layer, selected from [0.6, 0.8, 1.0]

### RFCN-DenseNet Search Space

Search space are as followings. 1) The blocks structure for DenseNet, default value is [[2, 3, 4],[3, 4, 5]]. 2) The growth rate of neuros in each block, default value is [8, 16, 32, 64, 128, 256, 512]. 3) The drop-out ratio of the first layer compared with the input layer, selected from [0.6, 0.8, 1.0]

### Res-VAE Search Space

Search space are as followings. 1) The number of layers, selected from [3, 4, 5]. 2) The drop-out ratio of the first layer compared with the input layer, selected from [0.6, 0.8, 1.0]. 3). The number of neuro in the first layer, selected from [4096, 2048, 1024, 512, 256, 128]. 4) The neuro number decay ratio of the next layer compared with previous layer, selected from [0.8, 0.6, 0.5].

### ENAS search

We improve the original ENAS^42^ methods in two ways to adjust to our framework. 1) We alter the original convolutional layer into fully-connected layer to adjust the input of modeling of genomic data. 2) We combine the output of different layers by concatenation operation. The number of layer in this mode should be fixed, default is 6. And the number of neuro in the first layer should be set in the JSON file (default is 2048). Search space are as followings. 1) The number of neurons in from the 2rd layer to the last layer, selected from [16, 32, 64, 128, 256, 512, 1024, 2048]. 2) The connection relationship between different layers.

### Feature importance estimation

We implement the SHAP^54^ package into AutoGenome to estimate the feature importance. When calling the function “autogenome.explain()”, AutoGenome take the best model and raw data as input for the SHAP module, with “GradientExplainer” mode specified. SHAP module automatically return the SHAP value of each feature for each sample. We also sum the absolute SHAP value of within each class and return the importance score of each feature for each class.

### Dataset preprocessing

All the dataset utilized in our study are public data. Data is first downloaded, followed by preprocessing procedures. AutoGenome automatically split the dataset into training, validation and test set with a proportion of 8:1:1. Details procedures of each case study are as followings:

### Experiment 1 – Pan-cancer classification

We downloaded gene expression profiles and somatic mutation profiles for pan-cancer patients’ tumor samples covering 28 human cancer types from The Cancer Genome Atlas (TCGA [https://gdac.broadinstitute.org/]) database. Log2-transformed Transcripts Per Million (TPM) was used to represent gene expression values. Somatic mutation profiles were extracted from TCGA mutation annotation files and represented by 0 or 1 indicating a gene is mutated or not in a sample. To avoid misleading classification, we filtered cancer types with a precise definition and removed four cancer types e.g. KIPAN, STAD, GBMLGG and COADREAD that included overlapped samples to other types, thus remained 24 cancer types for analysis. Then gene expression values of the 24 cancer types were processed by zero-one scaling in a gene-wise manner and input for model building.

### Experiment 2 - Mouse single cell

We utilized a dataset of single cell^50^ to perform a classification task. Expression matrix was downloaded as described in https://github.com/MarioniLab/EmbryoTimecourse2018. Due to the sample size is very large (more than 100,000 single cell), we randomly selected 10,000 single cell for downstream classification. The expression matrix we utilized contains 22018 genes for 10,000 single cells. These single cells belong to predefined 10 cell types, each have 1,000 cells. Expression matrix was subjected to 0-1 scale within feature before input to AutoGenome.

### Experiment 3

10X PBMC single-cell RNA-seq. The dataset was provided by 10X platform^76^. We downloaded the processed expression matrix and cell labels from (https://github.com/ttgump/scDeepCluster/tree/master/scRNA-seq%20data). This expression matrix contains 16,653 expressed genes for 4,271 single cells. These single cell belong to 8 predefined cell types, which corresponding to the colors in 3. Expression matrix was subjected to 0-1 scale within features, before input to AutoGenome.

### Reproducing the case studies

We will provide the source code and notebook examples to help researchers reproduce the case studies easily.

### Xgboost and Autokeras

We select “gbtree” in XGBoost as ‘booster’ to train the model, with the following parameters generated from a grid-like searched parameters, ‘objective’: “multi: softmax”, ‘gamma’: “0.1”, ‘max_depth’: “6”. We choose “MlpModule” in AutoKeras for genomics data modeling, with the parameter “loss = classification_loss” and “metric = Accuracy” and searched 10 hours for classification task

### PCA and T-SNE

“PCA” and “TSNE” function from “sklearn”^83^ were utilized for unsupervised learning for genomic data. After dimensionality reduction, the first two dimensions were utilized for further visualization. Then we calculate the pair-wise Euclidean distance of sample in the first two dimensions. As the raw data contain golden standard labels, so that we can calculate the Davies-Bouldin score to evaluate the performance of dimensionality reduction. “sklearn” was utilized to calculate the Davies-Bouldin score.

### Data Availability

All the data sets utilized in our study are public data. Pan-cancer classification is from TCGA. Single-cell classification data is from accessions: Atlas: E-MTAB-6967 and the processed data is downloaded following the instructions at https://github.com/MarioniLab/EmbryoTimecourse2018. 10X PBMC single-cell RNA-Seq was provided by 10X platform and we downloaded the processed expression matrix and cell labels from (https://github.com/ttgump/scDeepCluster/tree/master/scRNA-seq%20data).

### Software Availability

We will open the utilization of AutoGenome package to the public upon the acceptance of manuscript.

## Acknowledgements

We thank Prof. Dali Han and Prof. Guangdun Peng for critical reading and suggestions for revision.

## Author Contributions

N.Q. designed and conceived the project. W.H. and Z.X. implemented the AutoGenome python package and perform data experiment with the help from D.L. and C.X.. D.L., C.X. and L.Z. preprocess the data for experiment. J.Y. and W.F. contribute to the Hyper-parameter search and ENAS search module under the guidance of X.B. N.Q., D.L. and C.X. wrote the paper. X.B. D.H. and G.P. revised the manuscript. All authors read and approved the final manuscript.

