## Supplementary Figures for "AutoGenome: An AutoML Tool for Genomic Research"

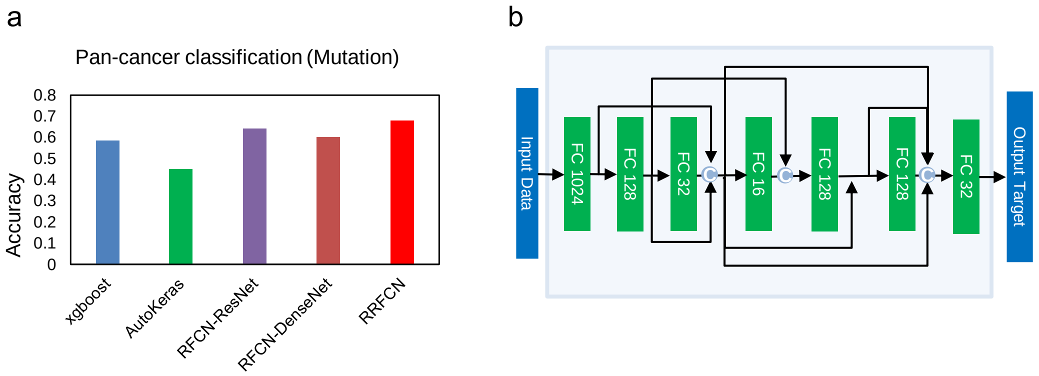


**Supplementary Figure 1. The pan-cancer classification using somatic mutation profiles.** a). Top-1 accuracy of XGBoost, AutoKeras, RFCN-ResNet, RFCN-DenseNet and RRFCN for the pan-cancer classification task using somatic mutation profiles. b). The best model architecture for pan-cancer classification using somatic mutation profiles.

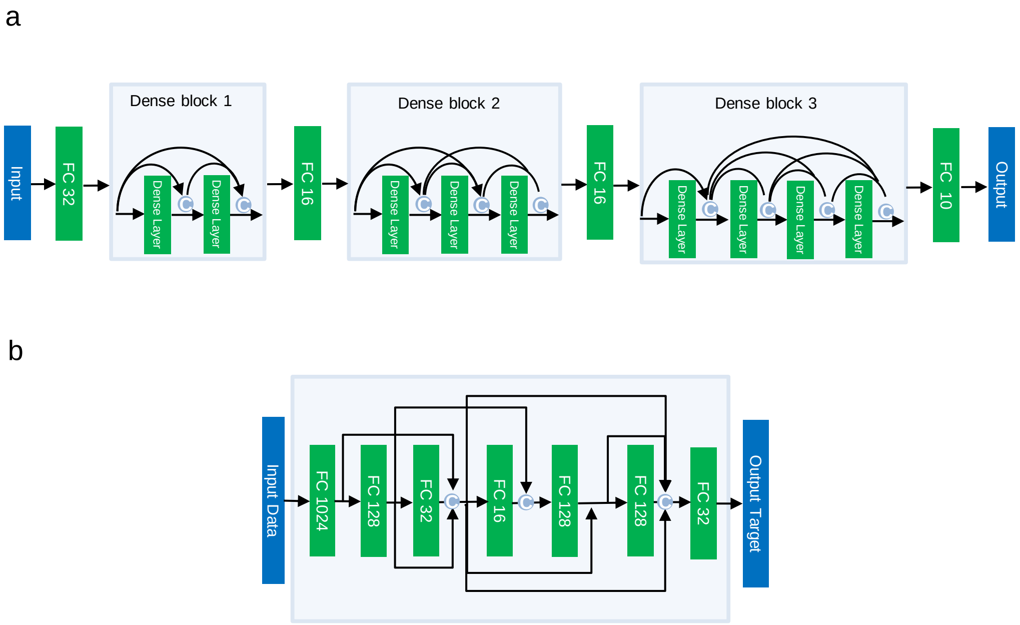


**Supplementary Figure 2. Model architectures for the single-cell classification task.**

a). RFCN-DenseNet and b). RRFCN architectures for the single-cell classification task.

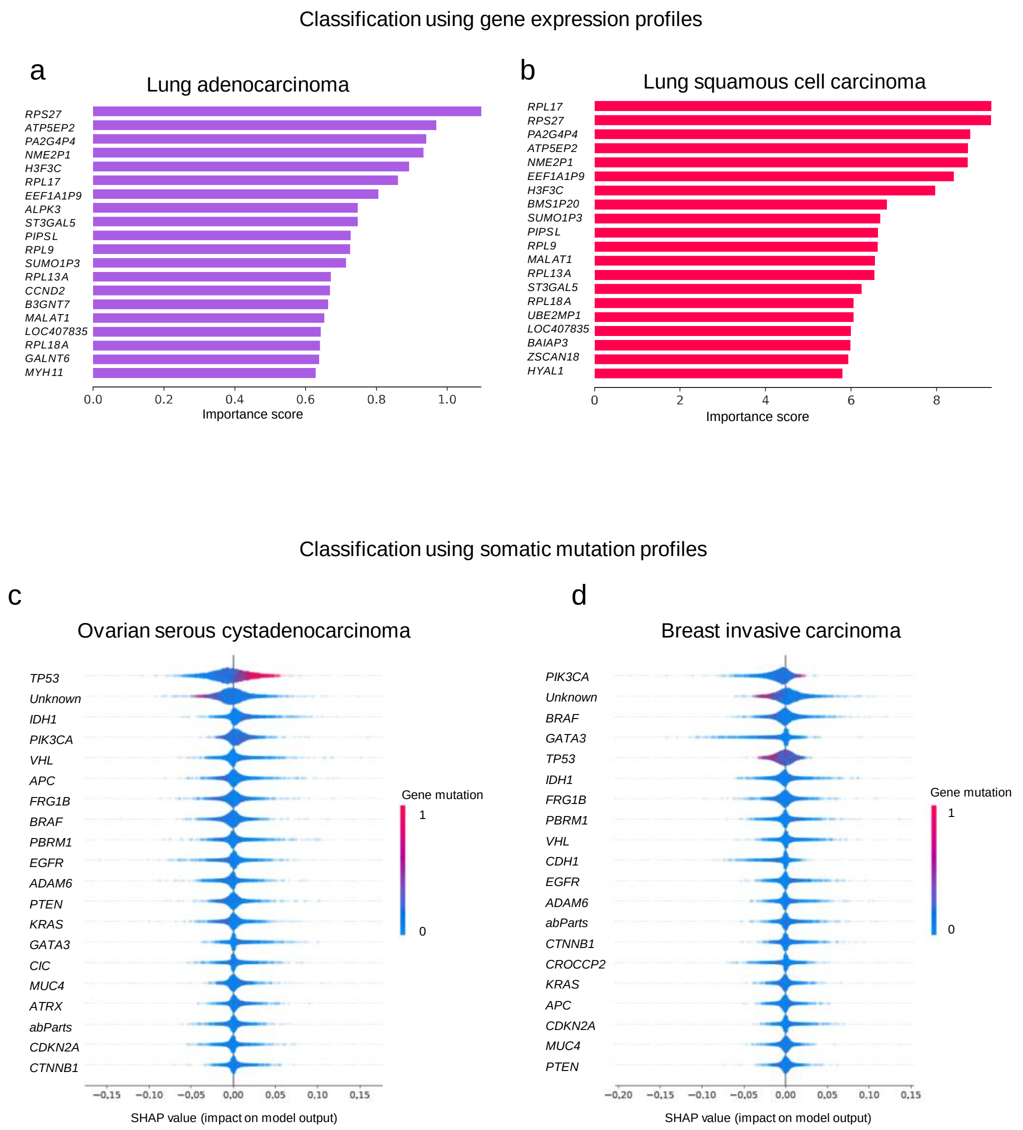


**Supplementary Figure 3. The feature importance analysis for the pan-cancer classification task.** Top-20 genes ranked by feature importance scores from gene expression based cancer type classification for a). Lung adenocarcinoma cervical and b). Lung squamous cell carcinoma, and from somatic mutation based cancer type classification for c). Ovarian serous cystadenocarcinoma and d). Breast invasive carcinoma. “Unknown” indicates regions that do not correspond to a gene. “abParts” indicates regions coding for highly variable portions of the immunoglobulin genes or for IGHV pseudogenes (named Ab-parts).

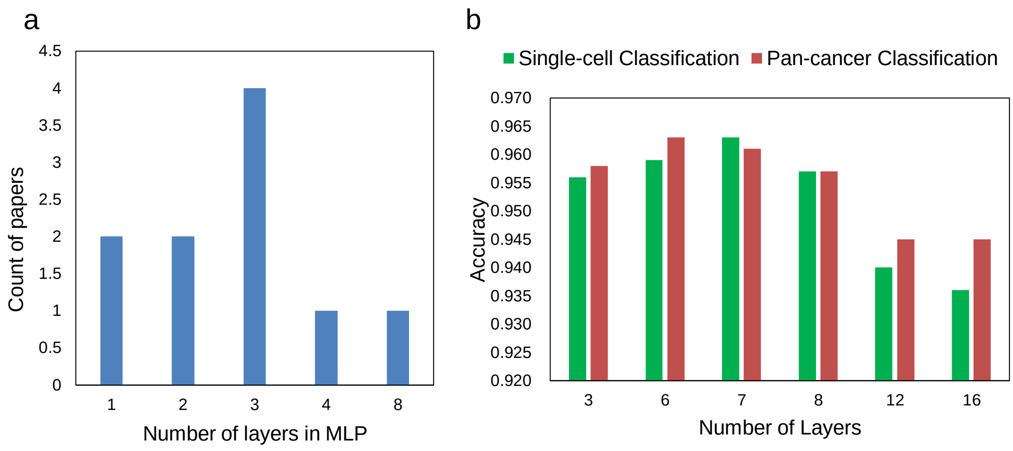


**Supplementary Figure 4. The number of hidden layers for MLP in published articles and its relationship to model accuracy in RRFCN.** a). Density plot of number of hidden layers for MLP in 10 published articles. b). The number of layers of models for the single-cell and pan-cancer classification tasks and corresponding top-1 accuracy.
